# Integration of Audiovisual Motion in Dorsolateral Prefrontal Cortical Neurons

**DOI:** 10.1101/2025.06.11.656227

**Authors:** Alireza Karimi, Rana Mozumder, Adriana Schoenhaut, Oscar Rausis, Mark Wallace, Ramnarayan Ramachandran, Christos Constantinidis

## Abstract

The dorsolateral prefrontal cortex is well recognized for its role in cognitive functions and activating action plans. In contrast, the properties of prefrontal neurons with respect to multisensory processing are less well studied. To address this question, we recorded single units from areas 8a and 46 of two female rhesus macaques while they were presented with visual, auditory, and audiovisual motion stimuli. The majority of dorsolateral prefrontal neurons responded to these sensory stimuli, with similar percentages of auditory-only, visual-only and audiovisual neurons. Approximately one third of responsive neurons exhibited significant super- or sub-additive interactions in response to the pairing of auditory and visual stimuli, revealing significant nonlinearities in their response profiles. Decoding motion signals from the population activity robustly differentiated multisensory from unisensory trials and also unisensory auditory and visual trials from each other. These results demonstrate that dorsolateral prefrontal neurons integrate auditory and visual motion signals, extending multisensory computations beyond sensory cortices into prefrontal circuits that support higher-order cognition.

**New & Noteworthy:** We recorded single neurons in macaque dorsolateral prefrontal cortex during visual, auditory, and audiovisual motion. Nearly half of responsive neurons were multisensory and a third displayed significant super- or sub-additive interactions, while ensemble activity reliably decoded stimulus modality. These findings provide the strongest evidence to date that DLPFC performs rapid, nonlinear audiovisual integration, extending multisensory computations beyond classical posterior regions into the prefrontal circuits that support cognition.

## 1 Introduction

Humans and other animal species navigate the world relying on a diverse array of sensory modalities. Each modality is tuned to specific forms of environmental energy, thus collectively increasing the probability of accurately detecting, discriminating, localizing, and identifying events or objects. In any given scenario, sensory information is often present from multiple modalities, thus making it challenging for the brain to decide what information should be segregated versus what information should be integrated or bound [1]. Since our perception of the natural world is a singular experience rather than a series of disparate perceptions derived from various senses, there must exist a mechanism for the integration of different sensory inputs. *Multisensory integration* is the process of combining inputs from multiple sensory modalities in order to modify (enhance or suppress) neural activity, with these neural changes presumed to reflect the brain substrates for the enhanced (or suppressed) the detection and identification of environmental stimuli [1, 2, 3]. In addition to facilitating behavior, these multisensory interactions are believed to strongly shape perceptual processes [4, 5, 6].

Audiovisual integration enables organisms to combine what they see and hear into a coherent percept, which is critical in many behavioral contexts. For example, watching someone speak while listening to their voice involves such integration, and consequently improves speech perception compared to hearing their voice alone, particularly in noisy environments [7, 8, 9, 10]. Numerous brain regions support audiovisual integration, spanning subcortical structures to high-level association cortices. Neurons in the ventrolateral prefrontal cortex (VLPFC) of monkeys have been shown to respond to auditory vocalizations and visual faces, and a subset of neurons integrate pairs of face-vocalizations [11, 12, 13, 14, 15].

The *dorsolateral prefrontal cortex* (DLPFC; areas 46 and 8a in monkeys; Fig. 1B) – long studied for its role in working memory and decision-making – has anatomical connections that could support multisensory convergence. Areas 46 and 8a receive convergent feed-forward projections from both the auditory belt/parabelt of the superior temporal gyrus and from dorsal-stream visual regions such as MT/MST, LIP, and posterior parietal cortex [16, 17, 18, 19, 20, 21, 22]. Moreover, reciprocal fronto-temporal loops also link DLPFC to the multisensory upper bank of the STS [12], parabelt of the superior temporal gyrus [23], and medial superior temporal area [24], providing a pathway for integrated audiovisual signals to reach prefrontal circuits. Prior studies in both humans and non-human primates have shown that neurons in DLPFC respond to visual, auditory, and audiovisual stimuli [25, 26, 27, 28]. These findings establish the presence of multisensory neurons in DLPFC. However, whether DLPFC supports true multisensory integration—as opposed to interdigitated auditory and visual representations—is a critical open question. Addressing this question has broad implications for understanding how sensory information is integrated within high-order cortical circuits responsible for goal-directed behavior and cognitive control.

**Figure 1:**
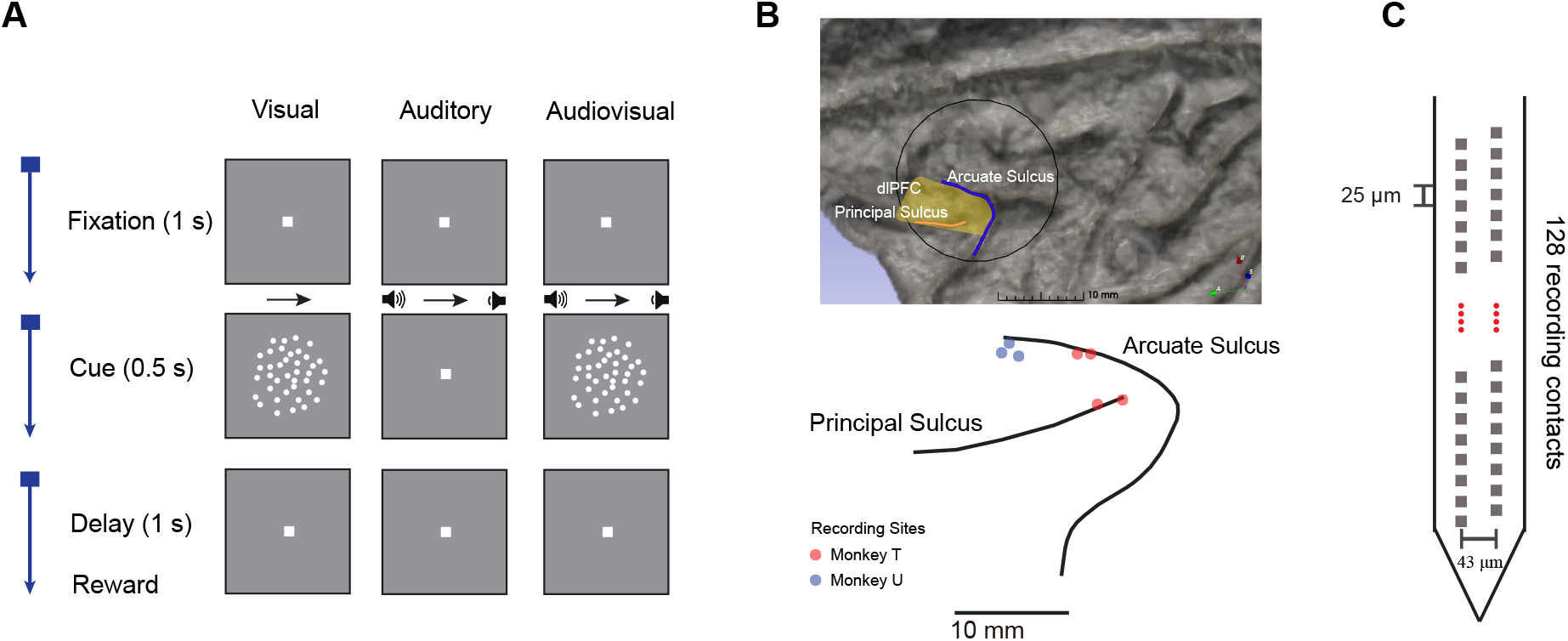
Task design, recording site and probe geometry. **A**. Each trial began with 1 s of fixation, followed by a 0.5 s stimulus (visual, auditory or audiovisual), and a 1 s post-stimulus delay. Visual stimuli were random-dot kinematograms (RDK) translating either leftward or rightward; auditory stimuli were broadband noise spatialised with inter-aural level differences to convey congruent motion (See §2.4 for details of stimulus); audiovisual trials presented both streams synchronously. Motion was presented with low or high coherence. **B**. Co-registered CT/MRI surface reconstruction of monkey’s brain. The recording chamber (black circle) targeted posterior and mid dorsolateral prefrontal cortex (yellow patch). Electrode penetration sites are specified with red circles for Monkey T (4 sessions) and with blue circles for Monkey U (3 sessions). **C**. Diagnostic Biochips laminar probe (128 sites in two staggered columns). Grey squares mark individual contacts.

Therefore, we were motivated to collect a large sample of neurons in the monkey DLPFC and identify their properties with respect to audiovisual stimulus pairings. Our experiment studied responses to random dot kinematogram (RDK) visual motion stimuli paired with apparent auditory motion sounds allowing us to probe how DLPFC neurons merge these signals to support audiovisual motion perception. We hypothesized that neurons in the DLPFC would integrate these audiovisual motion stimuli, thus exhibiting multisensory response enhancement or suppression relative to unisensory responses. The overarching hypothesis is that the DLPFC, traditionally associated with higher cognitive functions, actively integrates auditory and visual motion information to guide behavior.

## 2 Methods

### 2.1 Subjects

Two adult female rhesus macaques (*Macaca mulatta*; 8 years old) were used in this study. The monkeys were housed in shared environments, where they engaged in sensory interactions with their conspecifics. All surgical and experimental procedures were approved by the Vanderbilt University Medical Center Animal Care and Use Committee and were in accordance with the U.S. Public Health Service Policy on Humane Care and Use of Laboratory Animals and the National Research Council Guide for the Care and Use of Laboratory Animals.

### 2.2 Surgery and Neurophysiology

Each animal was implanted with a recording chamber (inner diameter 19 mm) in the left hemisphere centered on the dorsolateral prefrontal cortex. Cylinder placement and planned penetrations were verified by co-registering a post-operative CT with a structural MRI (aligned to the NIH Macaque template via @animal warper and visualized in 3D Slicer (see Fig. 1B).

Extracellular activity was recorded using multi-contact laminar probes from Diagnostic Biochips (Glenn Burnie, MD), featuring 128 recording sites arranged in two staggered columns with a horizontal spacing of 43.3 μm between columns and a vertical pitch of 25 μm (see Fig. 1C). Signals were acquired with the Open Ephys data acquisition system (OpenEphys system, Atlanta, GA) with a sampling rate of 30 kHz. Electrical signals recorded from the brain were amplified and band-pass filtered at 500–8000 Hz. Raw traces were common-average referenced and spike-sorted offline with Kilosort 2.5 (GPU-accelerated template matching with continuous drift correction; custom fork). Clusters were manually curated in Phy 2 and classified as single units if fewer than 0.5 % of inter-spike intervals fell below 1 ms, consistent with expected refractory-period violations.

### 2.3 Experimental Setup

The animals were seated in a primate chair with the head rigidly fixed inside a sound-attenuated booth. Visual stimuli were generated in MATLAB (Psychtoolbox 3; [29]) and presented on a 32-inch LCD monitor (Samsung QM32C, 1920 × 1080 pixels, 60 Hz; active area 70.5 × 40 cm) positioned 69 cm from the eyes.

Auditory stimuli were rendered at 48.828 kHz with PsychPortAudio and played through a pair of calibrated free-field loudspeakers mounted at ±27° azimuth.

Binocular eye position was tracked with an infrared video system (ISCAN ETL-200; 0.5° precision) whose analogue voltages were digitised at 1 kHz on a National Instruments DAQ and streamed to MATLAB in real time. TTL markers for stimulus onset, reward valve opening, and spike-sorting sync pulses were recorded on the same DAQ to align behavioural events with neural data.

All digital control lines (eye tracking, reward delivery, loudspeaker triggers) were routed through the same acquisition computer, ensuring sub-millisecond temporal correspondence between behavioural variables and the electrophysiological data described above.

### 2.4 Behavioral Task and Stimulus Set

The monkeys performed a *passive-fixation* paradigm—they were not required to make any choice or report about the stimulus, only to keep their gaze inside a 3° window around the central fixation point for the entire trial. Seven recording sessions were collected; the animals had no prior training on sensory-based tasks, only on fixation. An overview of stimulus conditions and recording geometry are summarised in Fig. 1.

Each trial consisted of four sequential epochs:

1. **Fixation acquisition** (max. 1 s). A 0.2° white point appeared at screen centre. The monkey had 1 s to bring its gaze inside a 3° radius window surrounding the point.
2. **Stimulus** (0.5 s). Stimuli from one of three modality conditions—visual (V), auditory or audiovisual (AV)—were presented while the fixation point remained visible.
3. **Delay** (1 s). No stimulus was shown; fixation had to be maintained.
4. **Inter-trial interval** (3 s). The fixation point was removed. A liquid reward was delivered only if fixation was held throughout the previous 3 steps; any break aborted the trial.

Visual stimuli were random-dot kinematograms (RDKs). Each RDK consisted of 50 white dots (0.1° diameter) presented within a circular aperture of 17° diameter. The dots were rendered at a luminance of 180 *±* 4 cd/m^2^ against a background luminance of 0.28 cd/m^2^, resulting in a Michelson contrast of 99.7 %. Visual motion was created by dots moved either leftward or rightward, at a constant speed of 40 deg s^*−*1^ throughout the trial. The motion coherence was manipulated by varying the proportion of dots moving in the target direction versus those moving randomly.

Auditory stimuli consisted of broadband white noise bursts spanning 0.1–24.4 kHz, sampled at 48.828 kHz. Stimuli were presented at 55.8±5.7 dB SPL (unweighted average over the band) via a pair of calibrated free-field speakers positioned at ±27° azimuth. Background noise in the booth, measured across the band, had a mean level of 23.8±6.1 dB SPL, yielding signal-to-noise ratios averaging 42.0±13.7 dB. The auditory motion signal was either leftward or rightward, embedded in partially correlated noise, and played through the speakers. The auditory stimuli consisted of four different components. First, individual white noise streams were presented through each speaker (100% amplitude; inter-signal correlation=0). Second, a combined white noise signal was presented through both speakers (100% amplitude, inter-signal correlation = 1). The fourth signal stream contained the apparent motion cue, in which the sound’s amplitude faded between the two speakers (decreasing the volume of one speaker while increasing the volume of the other over the stimulus duration) from 100% to 0% over the course of one trial (inter-signal-correlation = 0.5) to create binaural motion cues. This cross-fade produced an apparent angular speed of 108 deg s^*−*1^ (leftward or rightward) across the azimuthal plane. Coherence levels, representing the amplitude ratio of the faded signal to random jitters, were sampled from a log-spaced range. For the current paradigm, two representative levels were used: high (1.0) and low (0.5).

In audiovisual (AV) trials, auditory and visual motion stimuli were presented simultaneously in the same direction and with the same coherence, with their temporal onsets precisely synchronized.

The full factorial design produced **12 classes** (Table 1); each class specifies Modality, Direction (0 = left, 1 = right) and Coherence.

**Table 1:** Definition of the twelve stimulus classes used in every block.

| ClassID | Code | Description |
| --- | --- | --- |
| 1 | V 0 1 | Visual, left, high coherence |
| 2 | V 0 0.5 | Visual, left, low coherence |
| 3 | V 1 1 | Visual, right, high coherence |
| 4 | V 1 0.5 | Visual, right, low coherence |
| 5 | A 0 1 | Auditory, left, high coherence |
| 6 | A 0 0.5 | Auditory, left, low coherence |
| 7 | A 1 1 | Auditory, right, high coherence |
| 8 | A 1 0.5 | Auditory, right, low coherence |
| 9 | AV 0 1 | Audiovisual, left, high coherence |
| 10 | AV 0 0.5 | Audiovisual, left, low coherence |
| 11 | AV 1 1 | Audiovisual, right, high coherence |
| 12 | AV 1 0.5 | Audiovisual, right, low coherence |

Trials were organized into blocks, each continuing until the monkey successfully completed one trial from each class (12 correct trials per block). On each trial, a class was sampled uniformly at random without replacement from the remaining classes in the block. If the trial was performed correctly, the corresponding class was removed from the list; otherwise, it remained and could be re-selected. This ensured balanced condition counts across blocks while preserving a randomized, non-deterministic trial sequence.

### 2.5 Peri-Stimulus Time Histograms

Spike times were aligned to stimulus onset (*t* = 0 s; visual stimulus for V/AV, auditory stimulus for A). Firing rates were counted in 20 ms bins from *−*0.5 to +1.0 s. For population PSTHs (Fig. 6) each unit’s rates were *z*-scored to its baseline mean and standard deviation (−0.5 to 0 s) and then averaged; shaded regions denote *±*SEM. Gaussian smoothing (*σ* = 30 ms) was applied only to single-neuron exemplars (Fig. 3 and Fig. 4).

**Figure 2:**
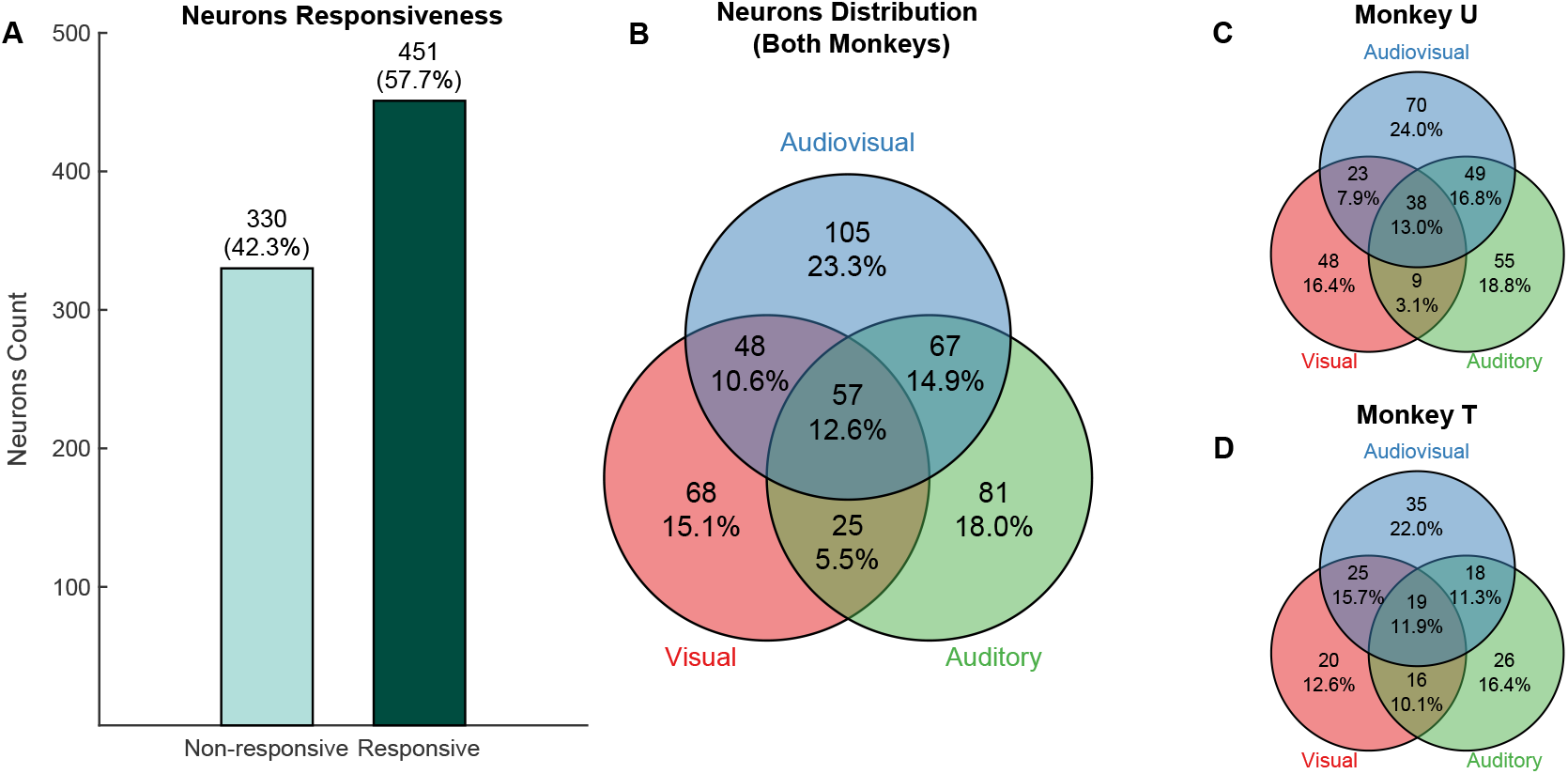
Sensory modality profile of dorsolateral PFC neurons. **A**. Number of responsive and non-responsive neurons. **B**. Modality specificity of the 451 responsive neurons pooled across both animals. **C**. Same analysis performed separately for each animal. For Animal U (*n* = 292 responsive units) and Animal T (*n* = 159 responsive units).

**Figure 3:**
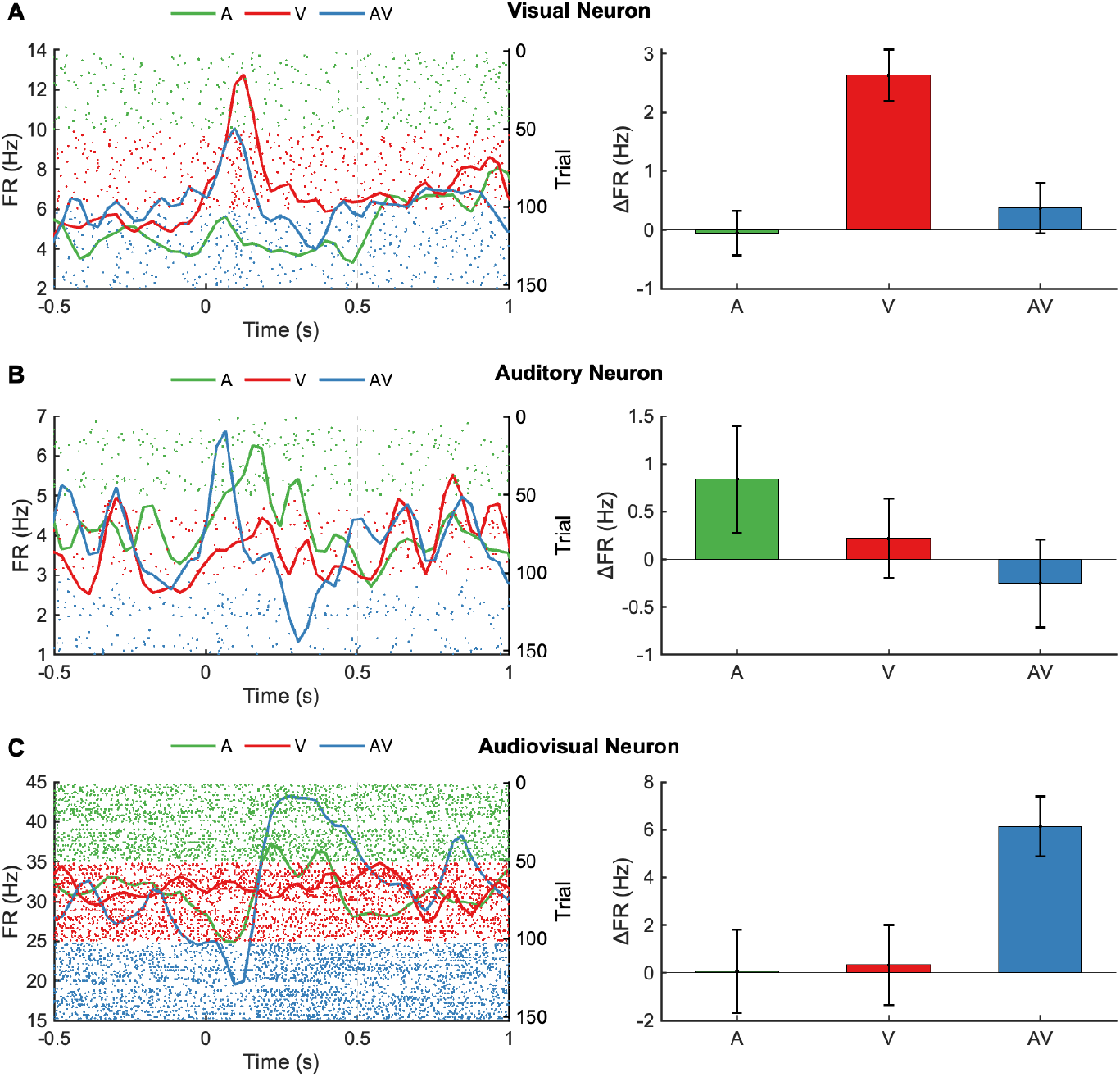
Modality-specific representative neurons. Each row shows a representative neuron classified as **A**. visual, **B**. auditory, or **C**. audiovisual. Left panels show spike density functions (SDFs) aligned to stimulus onset (time 0), separately for auditory (green), visual (red), and audiovisual (blue) conditions. Shaded gray vertical lines denote stimulus onset and the end of the post-stimulus analysis window (0.5 s). Right panels show the change in firing rate (ΔFR) from baseline to post-stimulus window for each condition. These example neurons are significantly responsive to one stimulus modality (*p <* 0.05) and not significantly responsive to the other two stimuli (*p ≥* 0.05).

**Figure 4:**
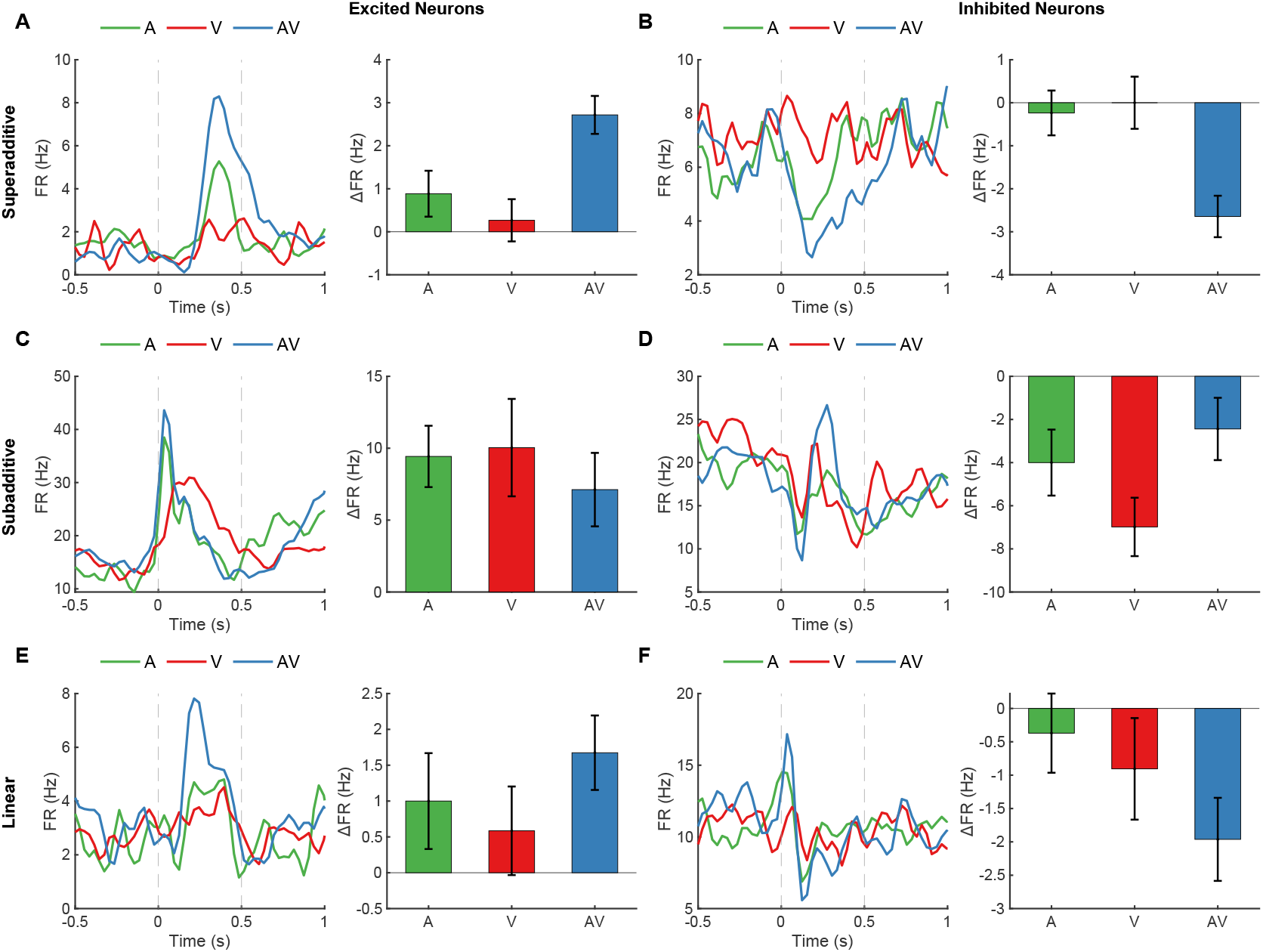
Representative examples of excited and inhibited neurons classified by multisensory integration profile. Rows separate functional classes defined from each unit’s audiovisual (AV) response relative to the arithmetic sum of its unimodal auditory (A) and visual (V) responses. In each panel, the left subplot shows the smoothed spike-density function (SDF), and the right subplot quantifies the firing rate change (ΔFR) during stimulation. Vertical dashed lines mark stimulus onset (0 s) and offset (0.5 s). Bars in right subplots represent mean ΔFR (post-stimulus FR 0–0.5 s minus baseline FR –0.5–0 s) with error bars denoting *±* SEM. **A**. Superadditive excited neuron (SI = 1.45, *p <* 0.005). **B**. Superadditive inhibited neuron (SI = 10.14, *p <* 0.001). **C**. Subadditive excited neuron (SI = –0.62, *p <* 0.004). **D**. Subadditive inhibited neuron (SI = –0.78, *p <* 0.001). **E**. Linear excited neuron (*p* = 0.447). **F**. Linear inhibited neuron (*p* = 0.354).

**Figure 5:**
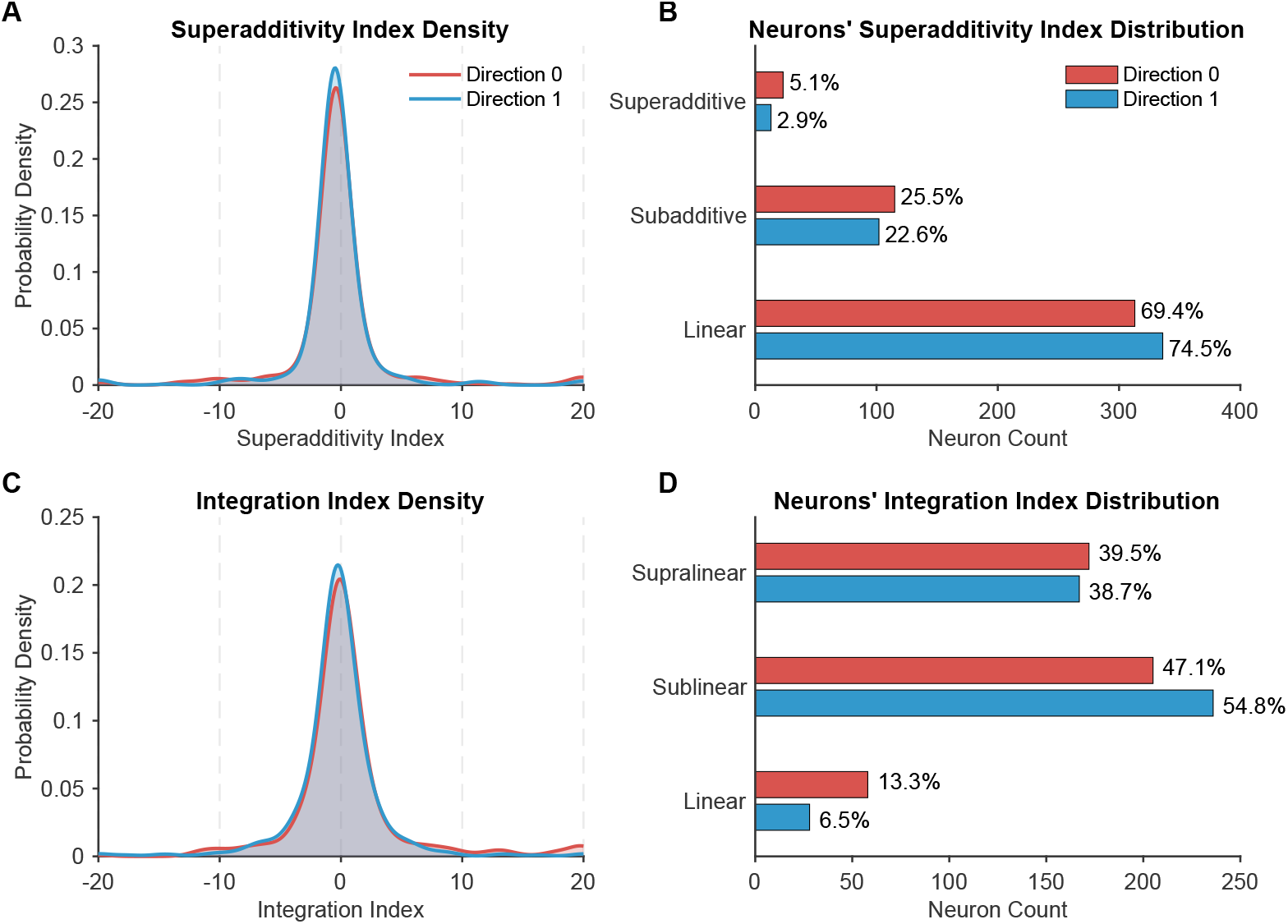
Superadditivity and integration index distributions among DLPFC neurons across motion directions. **A**. Kernel density estimates (winsorized at *±*20) of the superadditivity index (SI) for each motion direction. **B**. Grouped horizontal bars showing the number and percentage of neurons classified as superadditive (SI *>* 0 & *p <* 0.05), subadditive (SI *<* 0 & *p <* 0.05), or linear (*p ≥* 0.05). **C**. Kernel density estimates (winsorized at *±*20) of the integration index (II). **D**. Grouped bars showing counts and percentages of neurons classified as supralinear (II *>* 0 & *p <* 0.05), sublinear (II *<* 0 & *p <* 0.05), or linear (*p ≥* 0.05).

**Figure 6:**
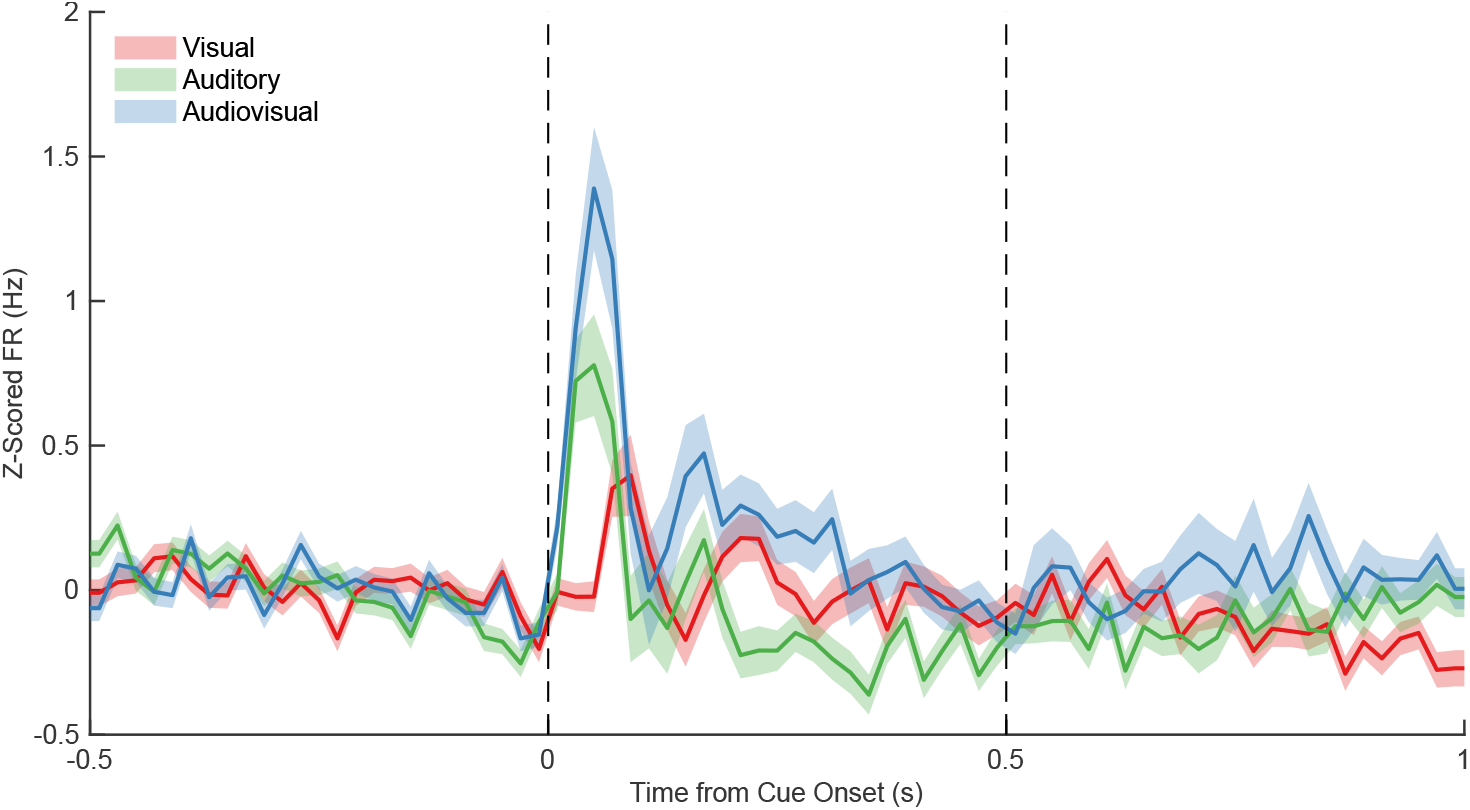
Population peristimulus time histogram of all responsive neurons. Firing rates (z-scored relative to the 500 ms pre-stimulus baseline) are plotted from –0.5 s before stimulus onset to 1.0 s after stimulus onset. Shaded bands represent *±* SEM across neurons. Vertical dashed lines denote stimulus onset (0 s) and offset (0.5 s).

### 2.6 Responsiveness and Selectivity

A unit was classified as *responsive* if its mean firing rate in the post-stimulus window (0 – 0.5s) differed from baseline (–0.5 – 0s) for at least one of the six modality-direction conditions (Wilcoxon signed-rank, *p <* 0.05; Fig.2A). Responsive units exhibited Modality, Direction, or Coherence *selectivity* when a two-sample ranksum test between the relevant stimulus groups reached significance (*p <* 0.05; Fig. 7). Being selective to a variable means that neurons have different levels of activity for different conditions of that variable.

**Figure 7:**
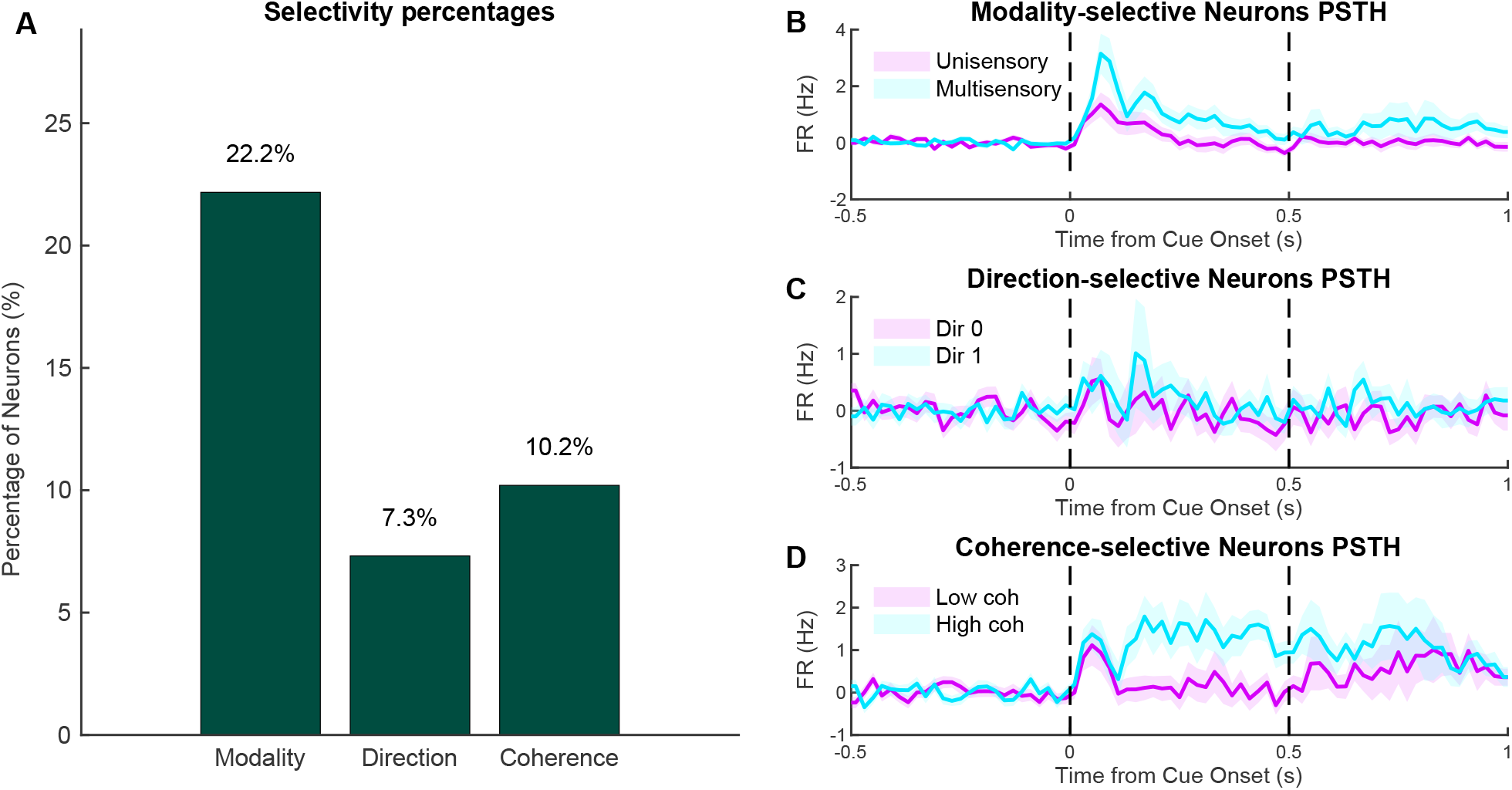
Population selectivity and evoked responses in dorsolateral PFC neurons. **A**. Percentage of responsive neurons showing significant selectivity for stimulus modality, direction, or coherence, based on firing rate differences across conditions (Wilcoxon rank-sum test, *p <* 0.05). **B–D**. Peristimulus time histograms (PSTHs) of z-scored firing rates from neurons selectively tuned for each feature. Shaded regions denote *±* SEM. Vertical dashed lines indicate stimulus onset (0 s) and stimulus offset (0.5 s).

### 2.7 Multisensory Integration Metrics

Baseline-corrected firing-rate changes were first obtained as

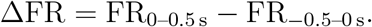

#### Superadditivity Index (SI)

The SI quantifies how strongly an audiovisual (AV) response exceeds—or falls short of—the linear sum of its unisensory components:

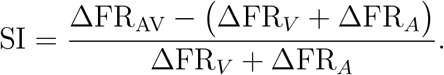

A positive value (SI *>* 0) indicates superadditive enhancement, SI *≈* 0 denotes an approximately linear combination, and SI *<* 0 reflects subadditivity.

#### Integration Index (II)

The II measures multisensory gain relative to the better single-modality response:

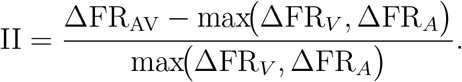

Here, II *>* 0 signals facilitation beyond the dominant unisensory response, II = 0 indicates no additional benefit, and II *<* 0 denotes suppression.

Indices were calculated separately for each motion direction. Significance was assessed by comparing AV trial ΔFR values with the trial-wise distribution of either (*V* +*A*) or max(*V, A*) using Wilcoxon rank-sum tests (*p <* 0.05; Fig. 5). Extreme values were Winsorized to *±*20 before kernel-density estimation.

### 2.8 Decoding Analysis

For each session, a linear support-vector machine (MATLAB `fitcsvm`, linear kernel) was trained to classify **(i)** Modality (unisensory vs. multisensory), **(ii)** Direction (0 vs. 1), and **(iii)** Coherence (low vs. high). The feature vector concatenated every neuron’s pre-stimulus firing rate (−0.5–0 s) with its post-minus-pre difference (0–0.5 s); features were *z*-scored per session. Performance was averaged over ten independent 70/30 hold-out splits (Fig. 8).

**Figure 8:**
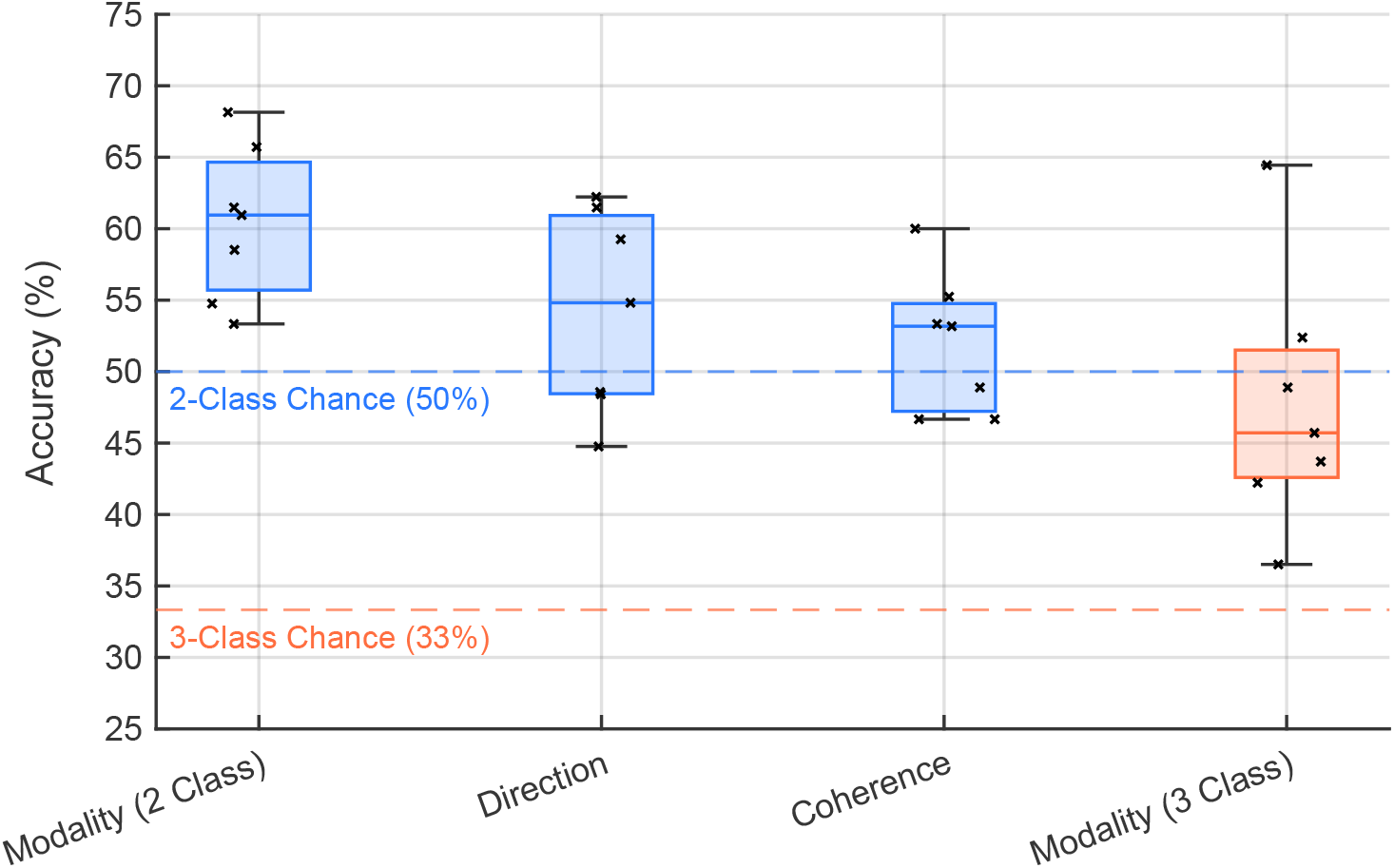
Decoding accuracy across sessions for Modality (2-class and 3-class), Direction, and Coherence. Box-and-whisker plots show classification accuracy across sessions using 10× hold-out validation with a linear SVM. Classifiers were trained to decode: (1) Modality (2-class) – multisensory vs. unisensory, (2) Direction – left vs. right, (3) Coherence – high vs. low, and (4) Modality (3-class) – visual, auditory, or audiovisual. Features included firing rate differences between post-stimulus (0–0.5 s) and pre-stimulus (–0.5–0 s) windows. Each box shows the interquartile range (IQR), with the median marked by a horizontal line and whiskers extending to 1.5× IQR. Individual session accuracies are overlaid as black crosses. Dashed horizontal lines mark the chance levels: 50 % for binary tasks and 33 % for the 3-class modality task. Mean accuracies across sessions were 60.9 % for Modality (2-class), 54.8 % for Direction, 53.2 % for Coherence, and 45.71 % for Modality (3-class).

### 2.9 Statistical Procedures

All statistical analyses were performed in MATLAB. Unless otherwise stated, tests were two-tailed with a nominal significance level of *α* = 0.05.

For every unit, a Wilcoxon signed-rank test compared the firing rate in the post-stimulus window (0–0.5 s) to the pre-stimulus baseline (−0.5–0 s) within each each of six modality-direction conditions (§2.6). This liberal criterion is designed exclusively to eliminate entirely unmodulated units. Any population proportions reported should be regarded as an upper limit; all subsequent analyses of selectivity and integration incorporate relevant statistical methodologies.

Modality, Direction and Coherence selectivity were assessed with two-sample Wilcoxon rank-sum tests that contrasted the relevant trial groups (§2.6).

Superadditivity Index (SI) and Integration Index (II) were computed per neuron and per direction (§2.7). Trial-wise AV ΔFR distributions were compared with either the (V + A) or max(V, A) surrogate distribution using rank-sum tests. Rank-sum test compared AV trials with the appropriate unisensory surrogate separately for Direction 0 and Direction 1. The two directions were analyzed and reported independently; therefore, no multiple-comparison adjustment was applied across them. To prevent extreme values from unduly influencing the kernel-density estimates shown in Fig. 5, index values were winsorised to the *±*20 range prior to density estimation.

Decoding accuracies obtained from 10 independent 70/30 hold-out splits were summarised with medians and inter-quartile ranges; box-and-whisker plots follow MATLAB defaults (box = IQR, whiskers = 1.5 × IQR).

## 3 Results

### 3.1 DLPFC neurons are broadly responsive to audiovisual motion stimuli

To determine the prevalence of stimulus-evoked activity in dorsolateral prefrontal cortex (DLPFC), we first compared baseline and post-stimulus firing for every isolated unit. A total of 781 well-isolated units were recorded from the dorsolateral prefrontal cortex (areas 8a and 46) of two macaques (n=520 from monkey U and n=261 from monkey T). A Wilcoxon signed-rank test (*α* = 0.05, two-tailed) compared each neuron’s firing rate during a pre-stimulus baseline window with its firing rate during an equal-length post-stimulus window (500 ms). A unit was classified as responsive if it exhibited a significant change for at least one of 6 modality-direction conditions (See Methods). This liberal screen is intended only to exclude completely unmodulated units. A total of 57.7 % (451 out of 781) neurons (292 out of 520 neurons = 56.1 % for monkey U and 159 out of 261 neurons = 60.92 % for monkey T) were deemed as responsive based on this criterion (Fig.2A).

Among responsive neurons, comparable fractions responded exclusively to visual (*V*; 15.1 %; Fig.3A), auditory (*A*; 18.0 %; Fig.3B), or audiovisual (*AV*; 23.3 %; Fig.3C) stimuli, while nearly one-third (43.6 %) responded to more than one modality condition (Fig.2B) and were classified as multisensory. Some neurons respond to multisensory stimuli without being responsive to unisensory motion stimuli (23.3 % AV-only neurons), whereas others respond to either unisensory or unisensory and multisensory stimuli. The roughly equal representation of each modality suggests that DLPFC receives convergent input from both visual and auditory motion pathways. The distribution of modality profiles was remarkably similar across the two animals (Fig. 2C), indicating that the basic pattern of sensory drive in DLPFC is robust across subjects.

### 3.2 Individual neurons exhibit multisensory computations

To illustrate how individual neurons combine audiovisual inputs, Fig. 4 shows representative neurons, classified by their multisensory integration profile. Rows in Fig. 4 separate units into units that show *superadditive* behavior (A-B), *subadditive* behavior (C-D), or *linear* behavior (E-F), based on the relationship between the AV response and the sum of the unisensory responses (see Methods). The left panels display smoothed spike–density functions (SDFs), and the right panels show the corresponding stimulus-evoked firing-rate change (ΔFR; post-stimulus minus baseline). Fig. 4A illustrates a neuron whose AV response exceeds the sum of its responses to V and A stimuli alone (superadditivity; SI = 1.45, *p <* 0.005). Fig. 4B shows a second superadditive example (SI = 10.14, *p <* 0.001), with a particularly large nonlinear enhancement. By contrast, Fig. 4C and Fig. 4D show neurons exhibiting subadditive interactions (SI = *−*0.62 and *−*0.78, *p <* 0.01 for both), in which the AV response is significantly smaller than the linear sum of the unisensory responses (i.e. suppressive interaction). Fig. 4E–F depict “linear” neurons whose AV response does not differ significantly from the sum of V and A responses (non-significant SI, *p >* 0.3).

Both excited and inhibited neurons in DLPFC can exhibit super-linear, sub-linear, or near-linear audiovisual responses, indicating that multisensory integration is distributed across cell types. This heterogeneity mirrors classical findings in cat and monkey superior colliculus, where multisensory stimuli can evoke marked enhancements or suppressions in neural activity [30, 1]. Importantly, a single AV stimulus does not elicit a uniform response across the population: some neurons amplify it, others suppress it, and many have responses that fall in between.

### 3.3 Heterogeneous nonlinear integration across the DLPFC population

A variety of integration profiles were observed across the population. Fig. 5 summarizes the distribution of multisensory interaction metrics for every responsive unit using the superadditivity index and integration index. Note that whereas the superadditive index compares the audiovisual response to the sum of the unisensory responses, the interactive index compares it to the stronger unisensory response. Using the superadditive index, about 30 % (30.6 % in one direction and 25.5 % in the other direction) of responsive neurons showed significant superadditive or subadditive interactions (SI *<* 0 or SI *>* 0; *p <* 0.05). The remaining 70 % (69.4 % in one direction and 74.5 % in the other direction) were statistically indistinguishable from linear summation (*p ≥* 0.05; Fig. 5B).

Using the integration index, roughly 40 % of the neurons exhibited multisensory responses that exceeded the strongest unisensory response (II *<* 0 or II *>* 0; *p <* 0.05; Fig. 5D). Importantly, direction-specific analyses revealed no systematic preference: the same neuron could be show superadditive responses for one direction of motion and linear or even suppressive responses for the opposite direction, underscoring the stimulus-specific heterogeneity of DLPFC multisensory computations.

### 3.4 Audiovisual stimuli elicit larger population responses

Peri-stimulus time-histograms pooled across all responsive neurons (Fig. 6) reveal that audiovisual firing diverged from baseline and reached a higher peak (mean z-score 1.4) than either unisensory distribution (visual: z=0.6; auditory: z=0.8). After stimulus offset (0.5 s), activity rapidly returned to near baseline in all conditions, indicating that these early responses are tightly tied to sensory drive. These dynamics show that integration occurs within the first 100 ms of cortical processing, well before decision-related activity typically appears in prefrontal neurons. This temporal distinction aligns with electrophysiological studies showing that sensory responses in PFC emerge rapidly—typically within 50–200 ms after stimulus onset [31, 32, 33, 34]. In contrast, decision-related signals, such as choice-predictive activity or evidence accumulation, arise later: approximately 200–300 ms after stimulus onset [35, 36]. Together, these findings support a temporal segregation between early multisensory integration and later cognitive computations such as decision formation in the PFC.

### 3.5 A substantial fraction of neurons selectively signal stimulus modality

We next asked whether single-neuron firing distinguishes stimulus modality independent of its motion parameters. Among the 451 responsive units, 22.2 % exhibited significant modality selectivity (unisensory or multisensory; *p <* 0.05; Fig. 7A). By contrast, only 7.3 % were direction-selective and 10.2 % were coherence-selective, indicating that stimulus type is the most salient single-neuron signal in this task. Modality-tuned neurons displayed classic cross-over PSTHs: firing was stronger for multisensory AV trials compared to unisensory A or V trials (Fig. 7B). Direction-selective cells (Fig. 7C) showed subtle difference in activity for left vs. right motion, while coherence selectivity (Fig. 7D) manifested more obvious differences in firing rate for coherence selective neurons.

### 3.6 Ensemble activity robustly decodes modality

Linear-SVM classifiers trained on population firing rates corroborated the single-neuron observations (Fig. 8). Across seven sessions, decoding accuracy for Modality in the binary contrast of multisensory versus unisensory trials averaged 60.9 %, reliably above the 50 % chance level (10*×* hold-out validation). When the task was expanded to a three-class discrimination among visual, auditory, and audiovisual trials, classification accuracy remained above chance, reaching a mean of 45.7 % compared with a 33 % chance level. Direction decoding was marginal (mean 54.8 %) and coherence decoding did not exceed chance (53.2 %). Because the classifier had access to both pre-stimulus baseline and post-stimulus changes, these results confirm that ensemble-level activity contains robust categorical encoding for sensory modality.

## 4 Discussion

### 4.1 Multisensory neurons are prevalent in dorsolateral prefrontal cortex

Multisensory processing was first described in subcortical and posterior cortical structures - including the superior colliculus (SC), where audiovisual interactions guide orienting behavior, and temporal-parietal regions such as the superior temporal sulcus (STS) and ventral intraparietal area (VIP), which combine visual, auditory and somatosensory information to encode objects near the body [30, 37, 38, 39]. Evidence indicates that the frontal lobe, and particularly the prefrontal cortex (PFC), also participates in multisensory processing, extending beyond its traditional role in cognitive control to encompass sensory integration. Although previous neurophysiological studies of the prefrontal cortex have focused disproportionately on visual stimuli, auditory responses have also been described [40, 41, 25, 26, 27, 28].

A handful of prior studies have described neurons responding to more than one sensory modality in PFC, however their prevalence varies considerably between studies, partly because of the small populations of neurons sampled. Watanabe (1992) recorded from pre-frontal and post-arcuate cortex and reported that 136 of 289 neurons were visually responsive, 13 responded to auditory stimuli, and 140 exhibited audiovisual responses [25]. Tanila et al. (1992) recorded from areas 46 and 9 in DLPFC during passive stimulus presentation and observed visual, auditory, and to a limited extent, somatosensory responsiveness. Out of 134 neurons, 50 responded to visual stimuli, 8 to auditory, and 5 to both [26]. Carlson et al. (1997) recorded from the monkey DLPFC and identified a small number of units responsive to both auditory and somatosensory stimuli, with 6 neurons responding to both modalities [28]. These studies established the convergence of information from multiple modalities within the same dorsolateral areas but they they did not systematically assess how neurons combined this information. The small number of neurons recorded in each study also made it difficult to assess the true prevalence of multisensory neurons and integrated responses.

Building on those observations, we recorded single units in dorsolateral PFC (areas 8a and 46) of two rhesus macaques during passive presentation of congruent auditory-visual motion stimuli. We observed a higher prevalence of auditory and audiovisual neurons in our sample compared to previous studies. More than half of the recorded neurons were classified as sensory responsive; similar percentages of neurons responded only to visual or auditory stimuli (15 and 18.0 % respectively), with almost half (44 %) exhibiting responses to the two modalities (Fig.2). Notably, a large fraction of these neurons responded to multisensory motion stimuli while remaining unresponsive to unisensory motion stimuli (23 % AV-only neurons). Consistent with the nature of the integrative responses reported in SC, STS, and VIP, a substantial proportion of DLPFC neurons (30 %) exhibited significant super- or sub-additive interactions, and 40 % responded more strongly to combined stimuli than to the most effective unisensory stimulus (Fig.5). Population decoding confirmed that ensemble activity carries a categorical code distinguishing AV, V, and A trials (Fig.8). Crucially, these nonlinear interactions emerged during the early sensory epoch—before the period of activity associated with decision-making or motor planning.

Our results are consisent with findings in the human prefrontal cortex. Clarke et al. (1995) used intracranial EEG in humans and reported 194 visual-responsive and 238 auditory-responsive sites in PFC [27]. Human fMRI studies further reveal a fine-scale mosaic of auditory and visual-biased domains in the lateral PFC that project to modality-matched posterior networks [42, 43]. In this scheme, modality-specific information is funneled through dedicated PFC microzones and then converges in adjacent domain general (multiple-demand) territory that implements task rules and decisions. It remains an open question whether a similar organization is present in the DLPFC of non-human primates.

### 4.2 Comparative landscape of brain regions implementing multi-sensory integration

A cardinal neurophysiological hallmark of multisensory integration is the presence of non-linear response interactions, where the neural response to a combined multisensory stimulus deviates from a simple linear summation of its responses to the constituent unisensory stimuli. Typically, a neuron is considered integrative if its response to a multisensory stimulus significantly differs from responses to the individual unisensory components under some criterion (e.g., greater than the maximal or sum of unisensory responses, or conversely, significantly suppressed relative to expectations) [30, 37, 44, 45, 11, 46, 1, 47]. Such effects are quantified with *multisensory indices* – for instance, a *superadditivity index* comparing the multisensory firing rate with the sum of unisensory rates, or an *integration index* comparing the multisensory response to the strongest single-modality response. These metrics, along with appropriate statistical tests, establish whether an area’s neurons actively synthesize inputs or simply respond in a modality-independent fashion.

In the present recordings from the monkey DLPFC, a significant portion of responsive neurons demonstrated such nonlinear integration. Specifically, based on the Superadditivity Index (SI), 25-30 % of neurons exhibited statistically significant (*p <* 0.05) superadditive or subadditive interactions. Superadditive responses (*SI >* 0) were rare, being observed in only 3-5 % of these neurons, while subadditive responses (*SI <* 0) were more prevalent, occurring in 22-25 % of neurons (Fig. 5A-D). The prevalence of these nonlinear interactions in DLPFC aligns with fundamental criteria used to identify brain areas actively involved in integrating sensory information. The observed heterogeneity, including both super- and sub-additive profiles, mirrors findings from other established multisensory regions, such as SC, STS, VIP, and VLPFC, though comparing multisensory metrics to other available studies of multisensory integration is challenging due to variability in methodologies and species.

A key distinction is that DLPFC in this study exhibited a greater tendency towards subadditivity contrasting with the more prevalent superadditivity seen in the SC [37, 5]. This divergence may reflect the differing computational roles of these areas, with the SC geared towards basic detection and orienting, and the DLPFC potentially engaged with more complex stimuli requiring efficient encoding rather than mere amplification. The Superior Temporal Sulcus (STS) and adjacent temporal regions are some of the major cortical sites for audiovisual convergence, particularly for complex and socially relevant stimuli such as faces, vocalizations, and biological motion. Studies indicate that a considerable fraction of STS neurons are modulated by audiovisual inputs; for example, Barraclough et al. (2005) reported that sound significantly modulated the visual response in 23 % of STS neurons with an equal split between enhancement and suppression [48]. In the parietal cortex, 56 % of the neurons recorded in the ventral intraparietal area (VIP) show significant modulation in response to stimuli from visual and tactile modalities. Of these, 41 % exhibited subadditivity and 15 % exhibited superadditivity [46].

In the ventrolateral prefrontal cortex (VLPFC) of macaques, multisensory integration is prominently observed, particularly in response to socially relevant stimuli such as faces and vocalizations. A study by Sugihara et al. (2006) recorded 387 neurons in the VLPFC and found that approximately 46 % (177 neurons) exhibited multisensory responses. This number is comparable with our findings. In VLPFC responses included both enhancements and suppressions when auditory and visual stimuli were combined. Notably, 10 % (40 out of 387 neurons) of VLPFC neurons were superadditive, and 28 % (110 out of 387 neurons) were subadditive [11].

### 4.3 Task and Stimulus Parameters Effects

An important point for the interpretation of the results is that the monkeys experienced the stimuli passively, without having to perform a judgment of motion direction, or any other stimulus-driven task. Learning and practicing tasks are well known to affect firing rate and neuronal correlations of prefrontal neurons [49, 50]. It is quite possible that the effects of integration would be more substantial had the monkeys been trained to perform a task that relied on a judgment about the multisensory stimulus. It is therefore quite notable that approximately two thirds of sensory-responsive DLPFC neurons were responsive to stimuli of both modalities. Similarly, it is noteworthy that one third of neurons exhibited significant super- or sub-additive interactions. Multisensory integration therefore appears to be a key feature of prefrontal processing, which provides the substrate for more complex interactions, following experience and learning [51].

Although neurons were categorized according to their sensory responsiveness and integrative profiles, it is important to point out that these are unlikely to be fixed categories. Stimuli were only manipulated within a limited dimensional space (i.e., three modalities, two directions, two coherences). Future work should explore parametric changes along these dimensions to fully reveal the computational operations performed by these neurons. As has been done in classic work in SC [52, 5, 53, 54], such experiments are likely to show that DLPFC neurons employ flexible operations based on stimulus characteristics. For example, a neuron originally labeled as subadditive may show additivity or superadditive interactions as it is probed along a specific stimulus dimension. Similarly, a neuron classified as visual may reveal itself as audiovisual when auditory feature space is further explored. Hence, the current study likely represents an underestimate of the true multisensory character of DLPFC, and only a glimpse into the likely rich landscape of multisensory integration in this cortical domain.

## Acknowledgements

We wish to thank Chrissy Suell and Kayla Yetman for valuable assistance during experiments and data collection. This research was supported by National Institutes of Health grant R01 EY017077.

## Disclosures

No conflicts of interest, financial or otherwise, are declared by the authors.

## Author Contributions

A.K. performed data analysis, generated figures, and wrote the manuscript. R.M. performed electrophysiological recordings. O.R. and A.S. designed and implemented the behavioral task. C.C. performed the surgical procedures. M.T.W., R.R., and C.C. initiated the research idea. R.R. and C.C. supervised the data analysis and contributed to writing the manuscript. All authors reviewed and approved the final version of the manuscript.

## Notes

### Competing Interest Statement

The authors have declared no competing interest.

